# Advax adjuvant formulations promote protective immunity against aerosol *Mycobacterium tuberculosis* in the absence of deleterious inflammation and reactogenicity

**DOI:** 10.1101/2020.10.01.323105

**Authors:** Diana H. Quan, Claudio Counoupas, Gayathri Nagalingam, Rachel Pinto, Nikolai Petrovsky, Warwick J. Britton, James A. Triccas

## Abstract

The development of safe and effective adjuvants is a critical goal of vaccine development programs. In this report, we defined the immunostimulatory profile and protective effect against aerosol *Mycobacterium tuberculosis* infection of vaccine formulations incorporating the semi-crystalline adjuvant δ-inulin (Advax). Advax formulated with CpG oligonucleotide and the QS-21 saponin (Advax^CpQS^) was the most effective combination, demonstrated by the capacity of CysVac2/Advax^CpQS^ to significantly reduce the bacterial burden in the lungs of *M. tuberculosis*-infected mice. CysVac2/Advax^CpQS^ protection was associated with rapid influx of neutrophils, macrophages and monocytes to the site of vaccination and the induction of antigen-specific IFN-γ^+^/IL-2^+^/TNF^+^ polyfunctional CD4^+^ T cells in the lung. When compared to the highly potent adjuvant combination of monophosphoryl lipid A and dimethyldioctadecylammonium bromide (MPL/DDA), Advax^CpQS^ imparted a similar level of protective efficacy yet without the profound stimulation of inflammatory cytokines and vaccination site ulceration observed with MPL/DDA. Addition of DDA to CysVac2/ Advax^CpQS^ further improved the protective effect of the vaccine, which correlated with increased polyfunctional CD4^+^ T cells in the lung but with no increase in vaccine reactogenicity. The data demonstrate that Advax formulations can decouple protective tuberculosis immunity from reactogenicity, making them ideal candidates for human application.

**Highlights:**

- Advax adjuvant formulations improve pulmonary protection against aerosol *Mycobacterium tuberculosis* infection
- Different combinations of adjuvant components markedly influence the level of protection observed
- Protection is associated with the rapid influx of myeloid cells to the site of vaccination and the induction of antigen-specific polyfunctional CD4^+^ T cells in the lung.
- Advax formulations abrogate vaccine-site ulceration and inflammatory cytokine production

## 1. Introduction

From prehistoric times to now, *Mycobacterium tuberculosis* has remained one of the most successful human pathogens. Tuberculosis (TB) is the leading cause of disease from a single infectious agent, causing an estimated 1.5 million deaths and infecting an estimated 10 million individuals in 2018 (1). The current vaccine, *Mycobacterium bovis* Bacillus Calmette–Guérin (BCG), is particularly effective at reducing the incidence of childhood TB, miliary TB and meningitis (2) but overall BCG efficacy in the lungs varies greatly depending on age of administration, localisation of TB infection, geographical area where vaccine is administered, previous exposure to various mycobacteria and current immune status (3). The negative global impact of BCG’s variable efficacy is profound because the lowest levels of protection occur in countries with the highest incidence of TB (4). WHO has estimated that this results in BCG only preventing 5% of all potentially vaccine-preventable deaths caused by TB (5).

In the search for a better TB vaccine, purified protein subunits are the safest and most refined design option and can be used in HIV-endemic areas, where it is not recommended to use live vaccine strategies. One important consideration is the fact that *M. tuberculosis* exists in two metabolic states, active or latent depending on whether infection is acute or chronic, and this can also impact vaccination strategies (6) since the antigen expression profile differs markedly between the two states (7, 8). Previous investigation of *M. tuberculosis* protein expression during chronic infection revealed strong induction of the sulfate assimilation pathway (SAP), the components of which proved to be highly immunomodulatory (9). This led to the development of CysVac2, a fusion protein consisting of early-stage immunodominant Ag85B antigen (Rv1886c) and the late-stage SAP protein, CysD (Rv1285)(10).

In combination with monophosphoryl lipid A (MPL) and dimethyldioctadecylammonium bromide (DDA), CysVac2 induces potent T helper 1 (Th1) and T helper 17 (Th17) responses, and confers a high level of protection in a mouse model of pulmonary TB infection (10). However, in order to improve the safety profile and reduce local reactogenicity events we have focussed on alternative adjuvant combinations, in particular the Advax family of adjuvants (11). Derived from δ-inulin, a plant storage carbohydrate, Advax has been shown to have a good tolerability and safety profile as an adjuvant and has been previously evaluated in vaccines against diseases such as hepatitis B, anthrax, listeria, SARS and influenza (12). We have previously demonstrated the immunogenicity and protective efficacy of adjuvanting CysVac2 with Advax in combination with a TLR9 agonistic CpG oligonucleotide (13). In this report we have further optimised this adjuvant formulation by evaluating the effects of additional adjuvant components that target different immune pathways. The saponin, QS-21, is a widely used, highly immunogenic adjuvant that induces strong antibody as well as cell-mediated immune responses, particularly Th1 and CD8^+^ T cell responses (14, 15). QS-21 has been used in clinical trials for malaria, influenza, HIV, hepatitis B, Alzheimer’s disease, and cancers (16), and is a key component of the M72/AS01 TB vaccine (17). The current study assessed combination adjuvants based on Advax to identify safe, well-tolerated regimens to protect against *M. tuberculosis* infection.

## 2. Materials and Methods

### 2.1. Bacterial Strains and Growth Conditions

*M. tuberculosis* H37Rv and *M. bovis* BCG Pasteur were grown at 37 °C in Middlebrook 7H9 medium (BD) supplemented with 0.5% glycerol, 0.02% Tyloxapol, and 10% albumin-dextrose-catalase (ADC) or on solid Middlebrook 7H11 medium (BD) supplemented with oleic acid–ADC.

### 2.2. Antigens and adjuvants

Protein antigens Ag85B (Rv1886c), CysD (Rv1285), CysVac2 were produced in recombinant form from *Eschericia coli* as described previously (10). Advax formulations, including CpG7909 and QS-21, were provided by Vaxine Pty Ltd (Adelaide, South Australia). DDA was purchased from Sigma Aldrich, Australia, and MPL was purchased from Invitrogen, USA.

### 2.3. Vaccination and infection of mice

Female C57BL/6 (6–8 weeks of age) were purchased from the Animal Resources Centre (Perth, Australia). Mice were maintained in specific pathogen-free condition and experiments were performed with the approval of the Sydney Local Health District Animal Welfare Committee (approval number 2013/047C) in accordance with relevant guidelines and regulations. Animals were randomly assigned to experimental groups. For protection experiments, mice were vaccinated subcutaneously (s.c.) at the base of the tail either once with 5 × 10^5^ CFU of BCG Pasteur (200 µl in PBS), or i.m. three times at two-week intervals with 3 µg of recombinant protein formulated in Advax (1 mg) in combination with other components: 10 μg CpG7909, 100 μg QS-21, 250 μg DDA, and/or 25 μg MPL. For intradermal (i.d) immunisation, mice were anaesthetised by intraperitoneal injection with ketamine/xylazine (80/10 μg/kg). Four microliters of protein and/or adjuvant (1 μg and/or 150 μg, respectively), adjuvant alone or PBS were injected i.d. into each ear under a surgical Leica M651 microscope (Leica, Wetzlar, Germany) using an ultrafine syringe (29G, BD Biosciences) as described by Li *et al*. (18).

For *M. tuberculosis* challenge experiments, mice were infected with *M. tuberculosis* H37Rv four weeks after their final vaccination via aerosol using a Middlebrook airborne infection apparatus (Glas-Col). Mice received an infective dose of approximately 100 viable bacilli. Four or 20 weeks later, lungs and spleens were harvested, homogenised and plated after serial dilution on supplemented Middlebrook 7H11 agar plates. Colony forming units (CFU) were determined approximately 3 weeks later and expressed as log_10_ CFU.

### 2.4. Cytokine production assays

Single cell splenocytes and lung cells suspensions were resuspended in buffered ammonium sulfate to lyse erythrocytes and then washed and resuspended in RPMI 1640 (Life Technologies) supplemented with 10% heat-inactivated foetal bovine serum (Scientifix, Cheltenham, Australia), 50 μM 2-mercaptoethanol (Sigma Aldrich, Australia), and 100 U ml^−1^ penicillin/streptomycin (Sigma Aldrich, Australia). Antigen specific IFN-γ producing cells were detected by ELISPOT assay as described previously (19). All antigens were used at a concentration of 10 µg ml^−1^. For cytokine bead array assays (CBA), cells were stimulated with antigens and supernatants collected after 72 hours. A cytometric bead-based multiplex assay kit (BD Biosciences) was used to measure the concentration of IFN-γ, TNF, IL-2, IL-17, IL-6 and IL-10 in accordance with the manufacturer’s instructions. Data were acquired on a BD LSR-Fortessa flow cytometer (BD) and then analysed using the FCAP Array Software (BD, USA).

### 2.5. Intracellular cytokine staining and flow cytometry

For intracellular cytokine staining, cells were stimulated for 3-4 hours in the presence of CysVac2 (10 µg ml^−1^) and then for up to 12 hours with brefeldin A (10 µg ml^−1^). Two million cells were incubated with anti-CD32/CD16 (eBioscience, San Diego, CA) in FACS wash buffer (PBS/2% FCS/0.1%) for 30 min to block Fc receptors, then washed and incubated for 30 min with either anti-CD3-PerCPCy5.5 (clone 145-2C11), anti-CD4-Alexafluor 700 (clone RM4-5), anti-CD8a-allophycocyanin (APC)-Cy7 (clone 53-6.7), or anti-CD44-fluorescein isothiocyanate (FITC) (clone IM7, BD). Fixable Blue Dead Cell Stain (Life Technologies) was added to allow dead cell discrimination. Cells were then fixed and permeabilized using the BD Cytofix/Cytoperm^™^ kit according to the manufacturer’s protocol.

Intracellular staining was performed using the following antibodies: anti-IFN-γ-phycoerythrin (PE)-Cy7 (clone XMG1.2), anti-TNF-APC (clone MP6-XT22, Biolegend, San Diego, CA), anti-IL-2-PE (clone JES6-5H4) (BD) or anti-IL-17A-Pacific Blue (clone TC11-18H10, Biolegend). For surface staining of ear samples preparations, cells were stained with anti-CD64-PE (clone X54-5/7.1.1), anti-MHCII-AF700 (clone M5/114.15.2), anti-CD45.2-BV510 (clone 104), anti-CD11c-PECy7 (clone HL3), anti CD11b-APC-Cy7 (clone M1/70), anti-CD326-APC (clone G8.8), anti-Ly6G-PB (clone 1A8), Ly6C-PerCPCy5.5 (clone AL-21).

All samples were acquired on a BD LSR-Fortessa flow cytometer (BD), and analysed using FlowJo^™^ analysis software (Treestar, Macintosh Version 9.8, Ashland, OR). A Boolean combination of gates was used to calculate the frequency of single-, double- and triple-positive CD3^+^CD4^+^ cell subsets. The gating strategy for intracellular cytokine staining is described in ref. (10).

### 2.6. Statistical Analysis

The significance of differences between experimental groups was evaluated by one- or two-way analysis of variance (ANOVA), with pairwise comparison of multi-grouped data sets achieved using Tukey or Dunnet post hoc tests.

## 3. Results

### 3.1. Advax^CpQS^/CysVac2 vaccination induces long-term protection against aerosol M. tuberculosis infection

We first examined how adjuvant formulations impacted on protective efficacy against *M. tuberculosis*. Mice were immunised with BCG or combinations of the CysVac2 antigen and Advax with CpG7909 (Advax^CpG^) or CpG plus QS-21 (Advax^CpQS^). CysVac2 formulated in MPL/DDA was used as a protective positive control formulation (10). Four weeks after immunisation, mice were challenged with aerosol *M. tuberculosis* and bacterial loads in the lungs determined. All adjuvant combinations and BCG protected mice against *M. tuberculosis* infection four weeks post-challenge when compared to unvaccinated mice, with the greatest effect observed with MPL-DDA/CysVac2 (Fig. 1A). At 20 weeks post-infection, the protective effect of BCG had waned, a finding we and others have observed previously (10). At this timepoint, only Advax^CpQS^/CysVac2 vaccination resulted in significant protection, reducing bacterial load in the lungs by approximately 0.52 log_10_ CFU compared to unvaccinated mice (Fig. 1B). Thus, Advax^CpQS^/CysVac2 was able to promote protective immunity against *M. tuberculosis* at both acute and chronic timepoints post-infection.

**Figure 1.**
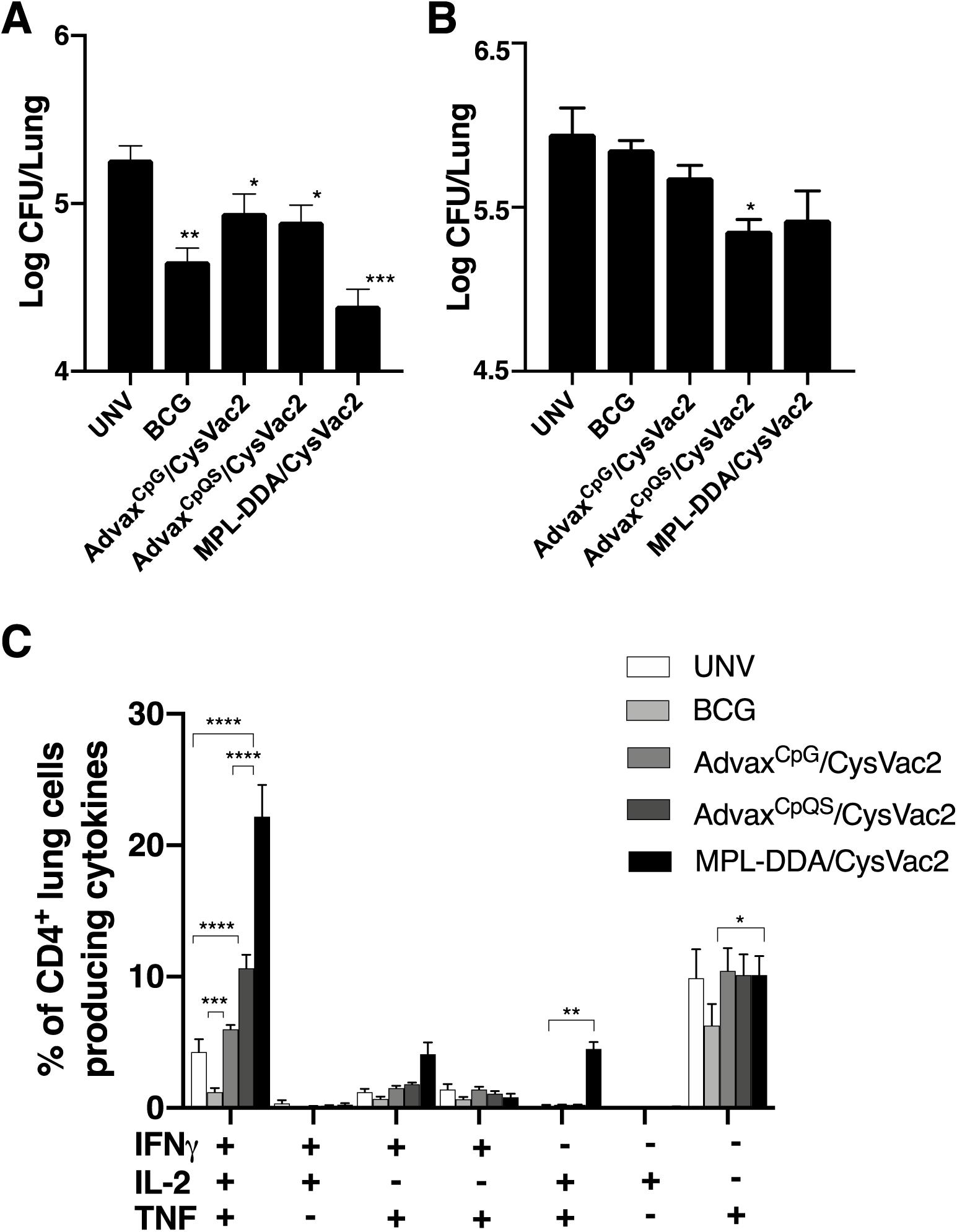
Protection against aerosol *M. tuberculosis* by Advax formulations. Female C57BL/6 mice (n=5-7 per group) were vaccinated once s.c. with BCG (5 ⨯ 10^5^ CFU), three times s.c. with 3 µg CysVac2 formulated in MPL-DDA, or three times i.m. with 3 µg CysVac2 formulated in Advax^CpG^ or Advax^CpQS^. Four weeks after vaccination, mice were challenged by aerosol infection with *M. tuberculosis* (∼100 CFU). Four weeks (A) and 20 weeks (B) post-challenge, the bacterial loads in the lung were determined and are shown as means ±SEM. Lung cells were restimulated *ex vivo* with CysVac2 in the presence of brefeldin A for the detection of intracellular IFN-γ, IL-2 or TNF via flow cytometry (C). The results are shown as means ±SEM for cytokine-producing CD4^+^ T cells. The significance of differences between the groups was determined by ANOVA (*P<0.05, **P<0.01, ***P<0.001, ****P<0.0001). Data are representative of two independent experiments.

To investigate vaccine-specific immune responses associated with the protection observed in Figure 1, mice were vaccinated and challenged as described above for Figure 1 and the generation of polyfunctional cytokine-secreting CD4^+^ T cells was examined four weeks post-infection. In Advax^CpG^/CysVac2-, Advax^CpQS^/CysVac2- and MPL-DDA/CysVac2-vaccinated mice, the vaccine-specific responses were dominated by IFN-γ^+^/IL-2^+^/TNF^+^ and TNF^+^ CD4^+^ T cell subsets (Fig. 2). BCG cytokine responses upon recall to CysVac2 were at background levels. While a similar pattern of responses was observed for all CysVac2 vaccinated groups, MPL-DDA/CysVac2 vaccination resulted in the highest proportion of polyfunctional CD4^+^ T cells (Fig. 2). Although a similar pattern was observed for Advax^CpQS^/CysVac2-vaccinated mice, the polyfunctional CD4^+^ T cell response to MPL-DDA/CysVac2 vaccination was significantly higher, and this was most apparent when examining the frequency of TNF^+^- containing subsets (Fig. 2). Thus, Advax is able to potentiate protective immunity when formulated with CysVac2, but without the heightened levels of inflammatory cytokine responses observed with MPL/DDA.

**Figure 2.**
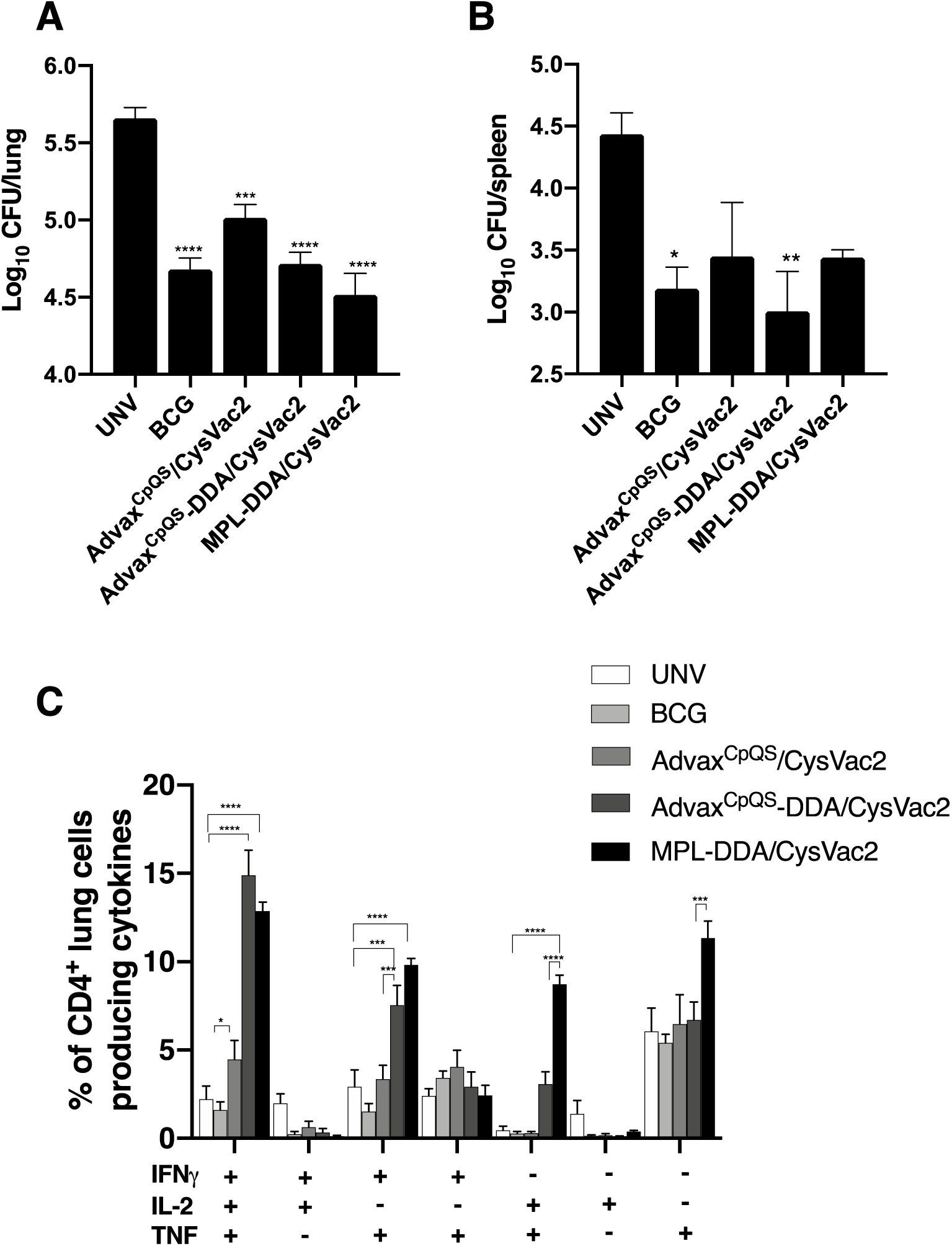
DDA augments Advax-mediated protective immunity against aerosol *M. tuberculosis*. C57BL/6 mice (n=5-7 per group) were vaccinated once s.c. with BCG (5 x 10^5^ CFU), three times s.c. with 3 µg CysVac2 formulated in MPL-DDA, or three times i.m. with 3 µg CysVac2 formulated in Advax^CpG^ or Advax^CpQS^. Four weeks after vaccination, mice were challenged by aerosol infection with *M. tuberculosis* (∼100 CFU). Four weeks post-challenge, the bacterial loads were determined in the lung (A) and the spleen (B), and are shown as mean ±SEM. Lung cells were restimulated *ex vivo* with CysVac2 in the presence of brefeldin A for the detection of intracellular IFN-γ, IL-2 or TNF via flow cytometry (C). The results are shown as mean ±SEM for cytokine-producing CD4^+^ T cells. The significance of differences between the groups was determined by ANOVA (*P<0.05, **P<0.01, ***P<0.001, ****P<0.0001). Data are representative of two independent experiments.

### 3.2. *DDA can augment Advax*^CpQS^*/CysVac2 protection against aerosol* M. tuberculosis *infection without enhancing vaccine reactogenicity*

Results in Figure 1 demonstrated that optimal protective immunity was observed with CysVac2 formulated in MPL/DDA, however this was associated with a strong inflammatory cytokine readout. We thus investigated if Advax^CpQS^/CysVac2 immunogenicity and protection could be improved by addition of DDA. Vaccination of mice with Advax^CpQS^-DDA/CysVac2 displayed protection equivalent to BCG and MPL-DDA/CysVac2 in the lung (Fig. 2A), and was the only CysVac2 formulation that significantly protected mice against systemic *M. tuberculosis* infection in the spleen, compared to unvaccinated mice (Fig. 2B). Analysis of cytokine-secreting CD4^+^ T cell subsets in the lung from vaccinated and challenged mice demonstrated that addition of DDA to Advax^CpQS^/CysVac2 markedly enhanced the triple positive IFNγ^+^/IL-2^+^/TNF^+^ population (Fig. 2C). However, the responses for other cytokine combinations were lower than those observed with MPL-DDA/CysVac2, particularly for double positive IFNγ^+^/TNF^+^ and single positive TNF^+^ producing CD4^+^ T cell subsets (Fig. 2B). When we further examined the level vaccine-specific cytokines in the supernatants of splenocytes restimulated with CysVac2, cytokine levels were highest in cells from MPL-DDA/CysVac2 mice, particularly for IL-2, TNF, IL-17A and IL-6. Splenocytes from Advax^CpQS^/CysVac2 released relatively low levels of cytokines compared to MPL-DDA/CysVac2, with only IFN-γ and TNF increased compared to background levels (Fig. 3). The addition of DDA to Advax^CpQS^-DDA/CysVac2 did not appreciably increase T cell cytokine responses. An increase in IL-17A was observed but this difference was not statistically significant (Fig. 3G). Finally, we examined reactogenicity of vaccines by determining the frequency of vaccine site ulceration across experiments. For all mice vaccinated with MPL/DDA, 12% developed ulceration (38 of 320 mice), with a minimum time to ulcer formation of 23 days. For all groups vaccinated with Advax, we observed no ulceration, irrespective of the adjuvant combinations used (0 of 221 mice). Thus overall, these results demonstrated that the Advax^CpQS^-DDA/CysVac2 combination can afford strong protective efficacy against both pulmonary and systemic *M. tuberculosis* infection but without deleterious inflammation and ulceration at the site of injection.

**Figure 3.**
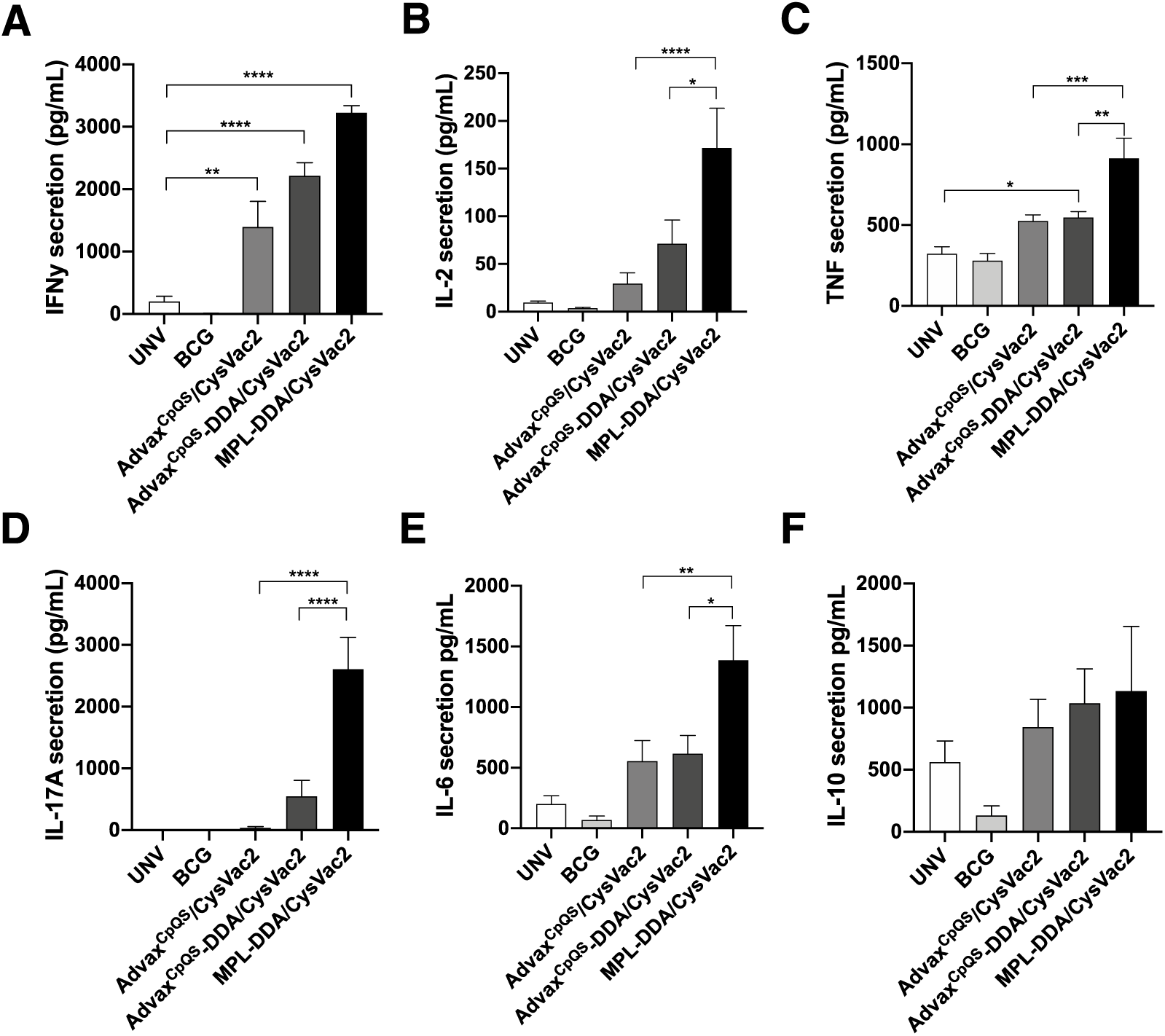
Reduced release of inflammatory cytokines after vaccination with Advax formulations. C57BL/6 mice (n = 4-7 per group) were vaccinated once s.c. with BCG (5 x 10^5^ CFU), three times s.c. with 3 µg CysVac2 formulated in MPL-DDA, or three times i.m. with 3 µg CysVac2 formulated in Advax^CpQS^ or Advax^CpQS^ with DDA. Four weeks after infection, spleen cells were harvested and restimulated *ex vivo* with CysVac2 over 72h. Cell supernatants were taken for the detection of secreted IFN-γ (A), TNF (B), IL-2 (C), IL-17A (D), IL-6 (E) or IL-10 (F) by CBA analysis in triplicate. The results are shown as mean ±SEM for mice in each group. The significance of differences between the groups was determined by ANOVA (*P<0.05, **P<0.01, ***P<0.001, ****P<0.0001). Data are representative of two independent experiments.

### 3.3. Cellular influx at site of injection after Advax/CysVac2 delivery

Early innate responses to vaccines shape the subsequent adaptive immune responses and are important for both persistence of memory and effector T cell functions. We used an i.d. delivery to the dermis of the ear to investigate cellular recruitment induced by CysVac2 vaccination (10), together with the gating strategy outlined in Supplementary Fig. 1. The day 2 response after vaccination with Advax^CpQS^/CysVac2 and Advax^CpQS^-DDA/CysVac2 vaccination was dominated by neutrophils, which were significantly elevated compared to unvaccinated or MPL-DDA/CysVac2 vaccinated mice (Fig 4A, 4B). NK cells, macrophages and, to a lesser extent, monocytes were elevated with Advax^CpQS^ formulations, however the addition of DDA reduced this response (Fig 4A, 4B). At day 4 post-vaccination, macrophages were the most abundant cell type at the vaccination site, and these were most apparent in mice vaccinated with Advax^CpQS^/CysVac2 and Advax^CpQS^-DDA/CysVac2 (Fig 4C). Neutrophils, monocytes and NK cells remained elevated in groups adjuvanted with Advax^CpQS^ compared to MPL/DDA, however for NK cells in particular, responses were markedly reduced for mice vaccinated with Advax^CpQS^-DDA/CysVac2. The numbers of T cells, DCs and B cells were not significantly different between groups (not shown). These results show that the protective effect of Advax-adjuvanted vaccines correlates with the early influx of distinct leukocyte populations to the site of injection.

**Figure 4.**
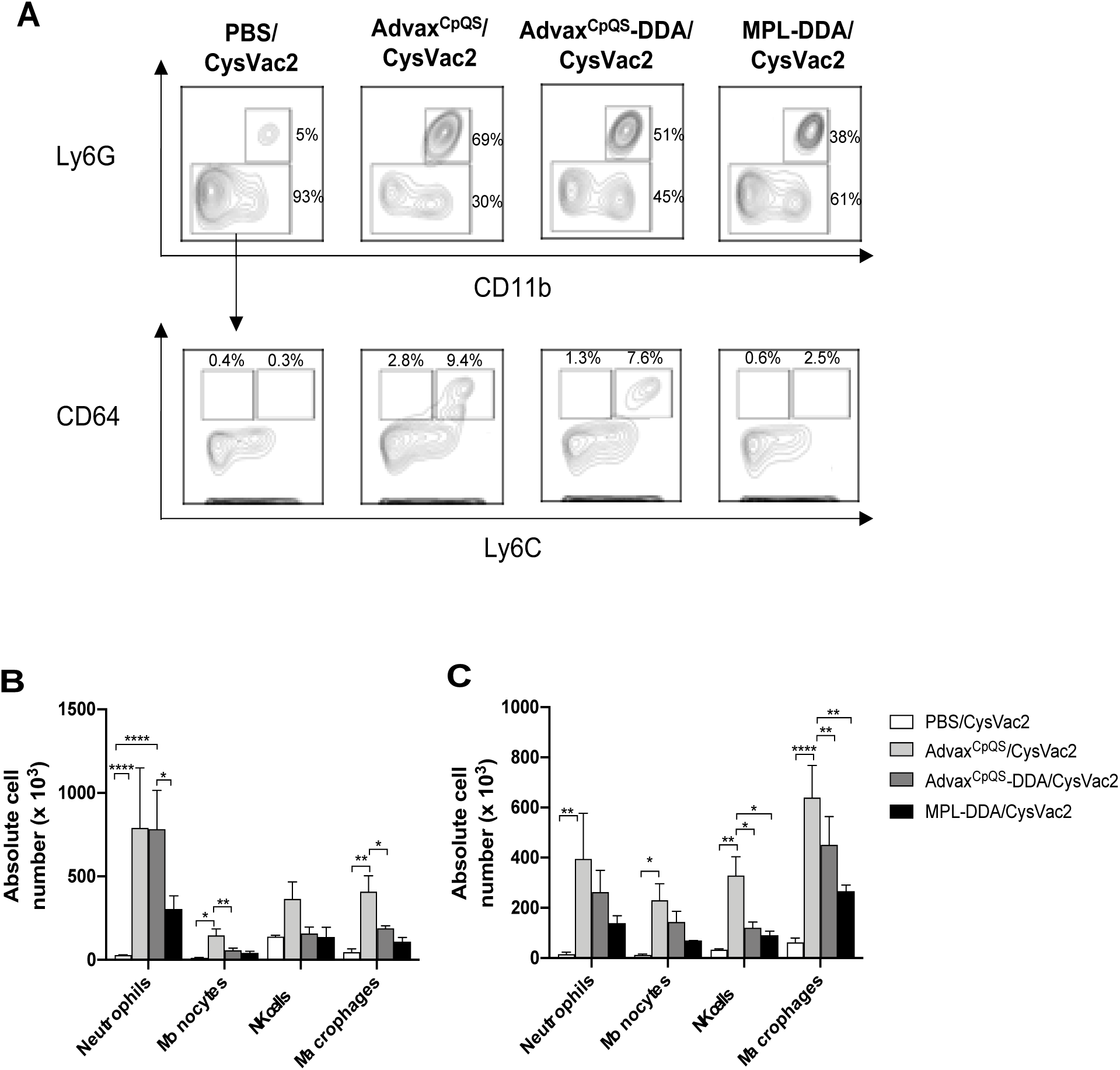
Leukocyte subsets recruited to the site of injection with adjuvanted CysVac2 formulations. C57BL/6 mice (n = 3) were i.d. injected in the ears with 1 µg CysVac2 formulated in PBS, Advax^CpQS^, Advax^CpQS^-DDA or MPL-DDA and the proportions and number of cell populations examined (A). Representative flow cytometry plots of neutrophil and monocyte/macrophage populations at day 2. The total number of neutrophils, monocytes, NK cells and macrophages are shown at two (B) and four (C) days post-injection and shown as mean ±SEM for the absolute numbers of the different cell types. The significance of differences between the groups was determined by ANOVA (*P<0.05, **P<0.01, ***P<0.001, ****P<0.0001).

## 4. Discussion

Successful vaccine development is a balance between effectively stimulating the immune system to mount a protective response while minimising adverse reactions upon administration. In this study, a fusion protein subunit vaccine consisting of CysD and Ag85B, termed CysVac2, was assessed in combination with various Advax adjuvant formulations. In accordance with previous results (10), vaccination with MPL-DDA/CysVac2 induced robust polyfunctional CD4^+^ T cell responses and protection against pulmonary *M. tuberculosis* infection in mice, on par with that afforded by BCG (Fig. 1A-B). However, MPL-DDA/CysVac2 vaccination was also associated with significant side-effects (injection site ulcer formation) in comparison to the Advax adjuvant formulations tested. This corresponded to increased inflammatory cytokine readouts in groups administered with MPL-DDA (Fig. 3C, E) and is consistent with the known reactogenicity of MPL-containing adjuvants (20). Furthermore, MPL has been shown to drive terminal differentiation of effector T cells and reduce protective capacity on antigen recall (48), suggesting MPL containing vaccines may not be ideal for vaccines seeking to provide long-term protection against mycobacterial infection. (21). We found that CysVac2 formulated with Advax combined with CpG and the saponin QS-21 afforded significant protection against pulmonary *M. tuberculosis* challenge, which was maintained at extended time-points post vaccination, unlike MPL-DDA (Fig. 1). QS-21 is also associated with dose-limiting side-effects when used alone (22), however we observed no adverse events with the Advax^CpQS^/CysVac2 vaccine in mice. This further supports the excellent safety profile of Advax-adjuvanted vaccines observed in pre-clinical and clinical studies (11).

Further optimisation of the Advax^CpQS^/CysVac2 formulation was achieved through incorporation of DDA, a potent immunostimulatory compound approved for use in human clinical trials (NCT00922363). DDA has long been recognised as an efficient adjuvant for TB subunit vaccines (23, 24) and is able to create a “depot” effect upon administration, which allows for undispersed antigen delivery to the draining lymph nodes to efficiently prime the adaptive immune response (25). The addition of DDA to Advax^CpQS^/CysVac2 resulted in a significantly greater induction of cytokine producing CD4^+^ T cells in the lung, although notably less TNF producing CD4^+^ T cells compared to MPL-DDA formulations (Fig. 2 and 3), highlighting the reduced inflammatory cytokine induction by Advax adjuvant combinations (26). In particular, the addition of DDA to Advax^CpQS^ increased induction of triple positive IFN-γ^+^/TNF^+^/IL-2^+^ producing CD4^+^ T cells in the lung (Fig. 2C). A greater proportion of double positive IFN-γ^+^/TNF^+^ and IL-2^+^/TNF^+^ CD4^+^ T cells were also observed in Advax^CpQS^-DDA/CysVac2-vaccinated mice (Fig. 3C), with the latter population having been shown to correlate with vaccine-induced protection against chronic *M. tuberculosis* infection (27).

When examining the effect of the different vaccine adjuvant formulations at the site of injection, Advax-containing vaccines resulted in a significant influx of macrophages and neutrophils to the dermis (Fig. 4A). Neutrophils are indicative of acute inflammation but also important in the activation of macrophages (28, 29) and direct killing of *M. tuberculosis* (30). The strong neutrophil response to Advax adjuvant may be due to the preferential phagocytosis of carbohydrates exhibited by neutrophils and enhanced by the structure of δ-inulin (31, 32). Neutrophils play a key role in macrophage recruitment (33) and this may explain the high levels of macrophages at the site of injection at 4 days post-immunisation in Advax groups (Fig. 4C). In contrast, MPL/DDA induced lower neutrophil and macrophage recruitment after CysVac2 vaccination. The greatest levels of NK recruitment to the vaccination were also observed in Advax^CpQS^/CysVac2-vaccinated mice (Fig. 4B-C). BCG vaccination has been demonstrated to induce trained immunity in NK cells and is postulated to play a role in the non-specific benefits of BCG vaccination such as decreased neonatal mortality (34). It is interesting to note that the addition of DDA to the Advax^CpQS^/CysVac2 formulation abrogated this NK response. This differs to the effect of MPL/DDA on NK cell recruitment when injected intraperitoneally (35) and may reflect difference in NK function at distinct anatomical sites. Nonetheless, as Advax^CpQS^-DDA/CysVac2 was the most protective formulation, this suggests that NK cells may not play a major role in the early induction of immunity after vaccination with this formulation.

In conclusion, this study demonstrates that potent protective immunity can be induced by the multistage CysVac2 fusion protein vaccine when administered with adjuvant combinations based on the non-inflammatory δ-inulin Advax platform. As Advax adjuvant formulations have been tested in both animals and humans without any major local or systemic reactogenicity (36-39), the vaccine formulations described in this study warrant further investigation to determine their potential to protect against *M. tuberculosis* in humans.

## Authorship contribution statement

D.Q., N.P., W.B. and J.T. conceived and designed the study. D.Q., C.C., G.N and R.P. performed the experiments. All authors analysed and interpreted the data. D.Q. and J.T. wrote the first draft of the manuscript. All authors reviewed and approved the final manuscript version.

## Declaration of Competing Interest

N.P. is an inventor on patents over Advax and has interests in Vaxine Pty Ltd, which owns interests in the Advax patents. W.B. and J.T. are inventors on patents over CysVac2. The authors declare that they have no known competing financial interests.

## Acknowledgement

This work was supported by a National Health and Medical Research Council (NHMRC) Project Grant (APP1102597), the NHMRC Centre of Research Excellence in Tuberculosis Control (APP1153493). We acknowledge the support of the European H2020 grant TBVAC2020 15 643381. The NSW Government provided support through its infrastructure grant to the Centenary Institute. D.Q. was supported by postgraduate scholarship funding from the NHMRC Centre of Research Excellence in Tuberculosis Control and an Australian Postgraduate Award. Development of Advax adjuvants was supported by funding from National Institute of Allergy and Infectious Diseases of the National Institutes of Health under Contracts, HHSN272201800044C, HHS-N272201400053C, HHS-N272200800039C and U01-AI061142.

