## Supplementary Figure 1 for "Advax adjuvant formulations promote protective immunity against aerosol *Mycobacterium tuberculosis* in the absence of deleterious inflammation and reactogenicity"

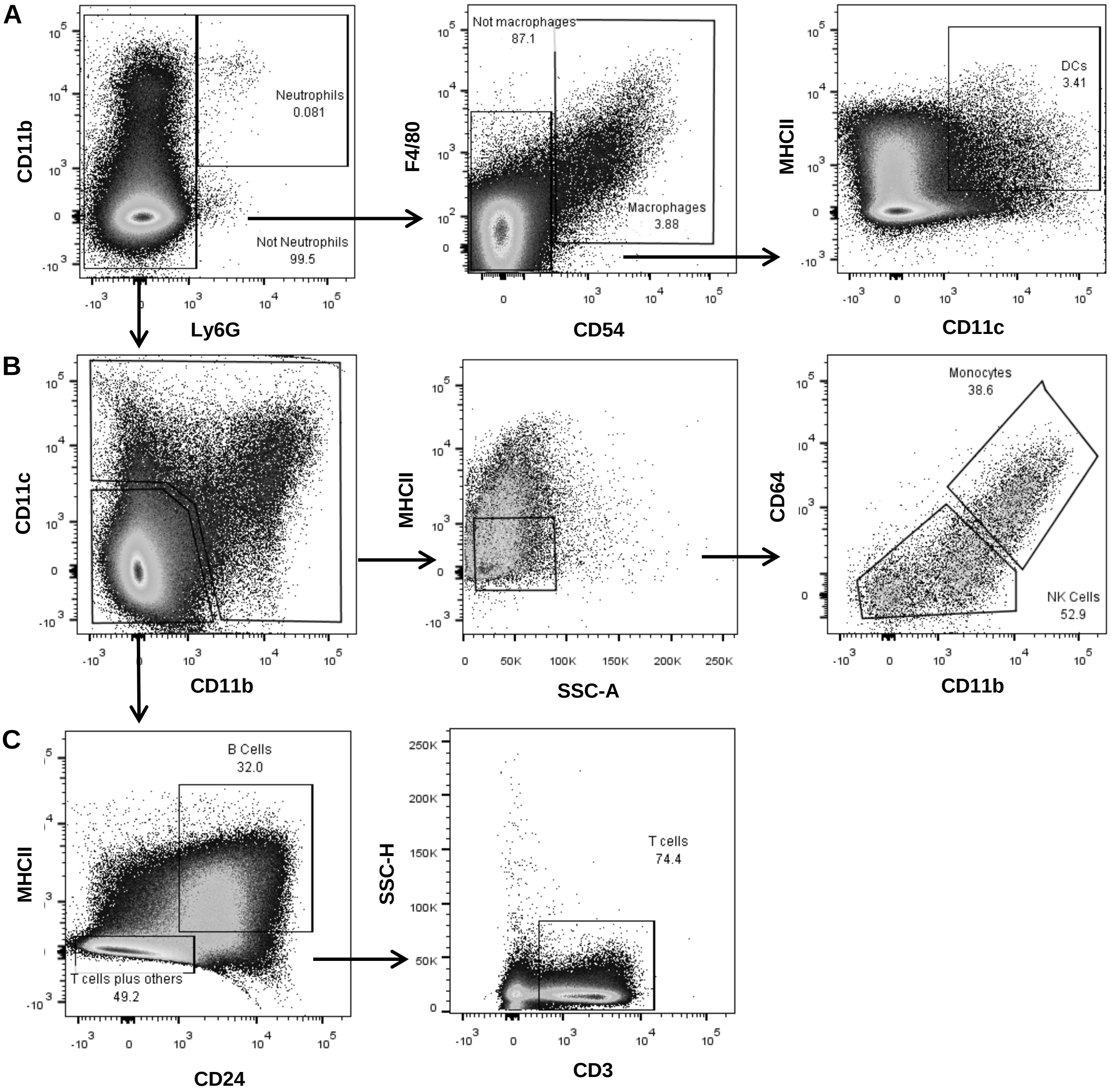


**Figure S1. Gating strategy for identification of leukocyte subsets.** After general gating to exclude debris, clusters and dead cells, this gating strategy was used to calculate proportions of neutrophils, macrophages and dendritic cells (A), the proportions of monocytes and natural killer cells (B) or the proportions of B cells and T cells (C). Method adapted from (Yu *et al.* 2016).
